# What is Antagonistic Pleiotropy?

**DOI:** 10.1101/321588

**Authors:** Josh Mitteldorf

**Affiliations:** Washington University School of Medicine, St Louis, MO, USA; National Institute for Biological Sciences, Beijing, China

## Abstract

Antagonistic Pleiotropy has been the dominant theory for evolution of aging since it was first proposed 60 years ago. Indeed, examples of pleiotropy have been observed, but there are also many examples of mutations that lead to longer lifespan without apparent cost. This poses a dilemma for the logic of the theory, which depends critically on the assumption that pleiotropy has imposed an inescapable precondition on evolution. Another interpretation is possible for the pleiotropy observed in nature. Natural selection may actually favor pleiotropy as an evolved adaptation. This is because the combination of high fertility and long lifespan is a temptation for individuals, but a danger for the health of populations. Predator populations that grow faster than their prey can recover are at risk of extinction. Once a sustainable mix of fertility and longevity has been established by multilevel natural selection, pleiotropy can help to assure that it is not lost. The population is free to shift from (high fertility/short lifespan) to (lower fertility/longer lifespan) as varying environmental conditions demand, without risking population overshoot and collapse. I describe herein experiments with an individual-based computer simulation in which pleiotropy evolves as a group-selected adaptation under a range of assumptions and in a broad swath of parameter space.

## Introduction

Antagonistic pleiotropy [1, 2] (AP) is currently the best-accepted explanation for the evolution of aging. The theory posits that there are genes (or evolved patterns of gene expression) for fertility and other essential elements of individual fitness that are so closely tied to loss of homeostasis and self-destruction late in life that natural selection has been compelled to accept the latter as a necessary cost of the former.

Senescence poses a dilemma for evolutionary theory because it clearly has a conserved genetic basis [3], despite its wholly negative impact on individual fitness. The advantage of the AP theory is that it resolves this dilemma within the context of uncontroversial selection mechanisms that are rooted in individual competition. The AP theory invokes no group selection. The disadvantage is that empirical evidence for pleiotropic links is spotty [4]. Many examples of pleiotropy have been discovered, but no evidence that they are unavoidable, or that they constrain natural selection in any meaningful way [5]. Worse, there are many known genes that accelerate aging for which no concomitant benefit has been identified. In his classic text on evolution of aging, Stearns [6] compiled a table of such genes across many species, and in the 26 intervening years, the number has grown beyond tabulation. Kenyon et al [7] describe cases of worms and other species as well in which disabling a gene that is prevalent in the wild type produces a healthy, long-lived mutant.

Spotty evidence has been interpreted as partial support [8], but I argue that it is incompatible with predictions of the AP theory [4]. Inconsistency of pleiotropy undermines the core logic of AP theory, because the theory depends critically on the assumption that the costs and benefits of pleiotropy are inseparable.

Contrary evidence includes both genetic and demographic examples.

There are (genetic) examples where senescence has evolved despite the fact that a clear, cost-free alternative is available. The most famous is fitness of the Trinidad guppy population as documented by Reznick [9], but there are many others [10–13]. Reznick found that guppies in a low predation environment had evolved lower fertility, shorter lifespan, and slower swimming speed, compared to a sister population just a few hundred meters distant in a high predation environment. Presumably, the lower fertility and shorter lifespan were necessary to keep population in check without the pressure of predation, but population regulation is a group benefit, incompatible with the framework of AP theory [14].

Demographic studies have generally not identified a “cost of reproduction.” Looking across the animal kingdom, Finch [15] concludes that reproduction often poses an immediate risk of death, but there is no evidence that fertility is linked to accelerated aging later on. Human demography finds no negative association between female fertility and longevity [16] and sometimes a small positive association is claimed [17, 18]. Some theorists argue that lack of evidence for a cost of reproduction should not be counted against the theory because natural selection has forged a compromise between fertility and longevity at the genetic level, not in any individual lifespan. The demographic evidence would be damning to the Disposable Soma theory, because it is based on metabolic tradeoffs, but not to other flavors of AP. But in all events, the many examples of cost-free genetic modifications that extend lifespan are a problem for AP theory.

For inconsistent pleiotropic links between fertility and senescence, another interpretation is possible. Within the context of adaptive theories of aging [4, 19–21], antagonistic pleiotropy may be an adaptive trait that preserves the long-term communal benefits of aging from being lost to the short-term power of individual selection. This perspective presumes that senescence has evolved by the long-term action of multilevel selection, as a compromise between the individual benefit of long lifespan and the communal risk of chaotic population dynamics. The combination of long lifespan with high fertility leads to population growth rates that are too rapid compared to regeneration time of the reservoir of food species. Population overshoot becomes severe, leading to chaotic population dynamics and extinction [4, 22]. Pleiotropy can be an evolved mechanism to avoid this Tragedy of the Commons [23]. Rapid, efficient individual selection is always pushing fertility higher and life spans longer, but if fertility and lifespan are inversely linked, the danger is avoided. A diversity of strategies can remain in the population with combinations of (long lifespan/low fertility) or (short lifespan/high fertility), but the population is protected from “cheaters” that might take over the population with an unsustainable combination of long lifespan and high fertility, pushing the population into overshoot and extinction. Keeping the product of lifespan and fertility in check is important for demographic stability of populations and ecosystems [4, 24, 25], and antagonistic pleiotropy as an evolved trait provides the benefit of suppressing the appearance of cheaters.

An objection might be raised that pleiotropy is everywhere in the genome, and no evolutionary mechanism is required to explain the provenance of pleiotropy in this particular case. But universal pleiotropy derives from nature’s economy, the re-use of old tools for new tasks. Pleiotropy results when evolution proceeds along a path of least resistance. *Antagonistic* pleiotropy demands an evolutionary explanation because it is not the dual use of a single trait, but rather the compelled acceptance of a linkage that is costly to individual fitness.

From the framework of conventional population genetics, another objection arises: that “natural selection has no foresight”. It seems like teleological thinking to imagine that natural selection might be so wise as to protect senescence against the force of short-term selection because it poses a risk of extinction from population overshoot.

To this, I respond with (1) an uncontroversial example of a parallel case in which natural selection has created just this sort of protection against short-term advantage, and (2) a simple computational model, including group selection and individual selection in a natural balance, in which it can be seen that pleiotropy evolves handily as an adaptation.

## Precedents

The uncontroversial example is evolution of the linkage between sex and reproduction which appeared around the time of the Cambrian explosion. If sharing genes were optional, there would be strong individual selection to avoid sharing genes. Gene-sharing wastes time and energy. For the best-adapted combinations of genes, there is a cost to sharing genes with the competition. Sex became linked to reproduction precisely to make it obligatory. In most multicelled animals and plants, there are powerful genetic and anatomical barriers that prevent evolutionary reversion to asexual reproduction. The linkage between reproduction and sex is so familiar to us that we may forget that the linkage itself must have adaptive value, else it could not have evolved.

Sex in the guise of conjugation first evolved in single-celled *protoctista*, and for hundreds of millions of years, sex was an activity independent of reproduction [26]. Reproduction took place via cloning, and sex took the form of conjugation. Sex had no adaptive value for the individual, and the first mechanism that evolved to enforce the sharing of genes was cellular senescence, an early mode of programmed aging. In ciliates, telomerase remains sequestered through the life cycle, hence telomeres are permitted to shorten during mitosis. Telomerase is released only during conjugation, requiring that every lineage must submit to conjugation periodically or it will succumb to telomere exhaustion and cellular senescence.

With the appearance of multi-celled life, this method of making sex obligatory was no longer available. Quite remarkably [27], sex become anatomically bound to reproduction, enforcing the imperative to share genes in every life cycle. Among evolutionary theorists, the purpose of sex itself has been debated roundly for decades; but for the link between sex and reproduction, there can be only one conceivable purpose, and that is to enforce gene sharing, by effectively blocking the devolution to an asexual mode of reproduction. The reason that sex became a prerequisite to reproduction is that the selective advantage of sex accrues only to the deme, and the cost is borne by individuals. Without this tight linkage between sex and reproduction, sex would easily be lost to the efficient operation of individual selection and the community would lose its diversity and its capacity to adapt in a changing environment.

Thus the evolution of obligate sexual reproduction provides an antecedent for exactly the mechanism that I propose is responsible for the evolution of pleiotropy. Both sex and aging have group advantages and individual costs. Both are in danger of disappearing unless they are protected from the quick and efficient operation of individual selection. Sex is protected from individual selection by dividing the means of reproduction between two individuals, so that in order to replicate, every individual must share genes. Aging is protected from individual selection by many and various modes of genetic pleiotropy, linking longevity to substantial fitness costs.

Like sex, antagonistic pleiotropy is an evolvability trait, which is to say it has no effect on fitness in the here and now, but modulates the dynamic of natural selection over evolutionary time. The evolution of evolvability is not accordant with standard models of population genetics [28], but we know nevertheless that it has occurred [29, 30]. In addition to sex, common evolvability traits include differential mutation rates, the hierarchical structure of the genome, the maintenance of diversity, aging, and the genetic code itself [29, 31, 32]. Whether or not we accept proposed mechanisms for selection of evolvability adaptations, we must respect the fact that somehow evolvability has evolved.

## The Delayed Logistic Model

The computational model for the evolutionary emergence of AP is based on a metapopulation of local sites, each of which is governed by delayed logistic dynamics [22, 24, 33]. The logistic equation is a venerable model for behavior of populations subject to a fixed carrying capacity. 
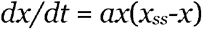

Solutions of the logistic differential equation are always stable, with population x restoring smoothly and efficiently to the steady-state level x_ss_. But in the real world, feedback concerning the sustainable steady-state level of population is not immediate. For animals, there is always a reservoir of food in the environment. In the long run, the reservoir is renewable, but in the short run the food reservoir can be substantially depleted by unsustainable consumption. As long as food can be found, reproduction may proceed unimpeded, oblivious to the impending food shortage. The population may continue to overshoot its steady-state level until severe food shortage causes a massive die-off. Thus, food availability provides a basis for logistic population dynamics in which the restorative term in the equation is generally delayed in time. In contrast to the differential form of the logistic equation, solutions to the logistic equation with time delay can be either stable, fluctuating, or chaotic.

In the early history of chaos theory, the logistic difference equation with discrete time was the first equation used to demonstrate chaotic dynamics [34, 35]. The choice between smooth behavior and oscillating or chaotic behavior is predicted by the chaos parameter, defined as the product of the discrete time step Δt and the (logarithmic) population growth rate. If the chaos parameter is small, the population restores smoothly to its steady-state level. If the parameter is more than 2, there is some oscillation. As the chaos parameter approaches 3, the population curve behaves chaotically. For values >3, the population overshoots, then goes extinct in a single cycle. (Fig. 1)

**Figure 1.**
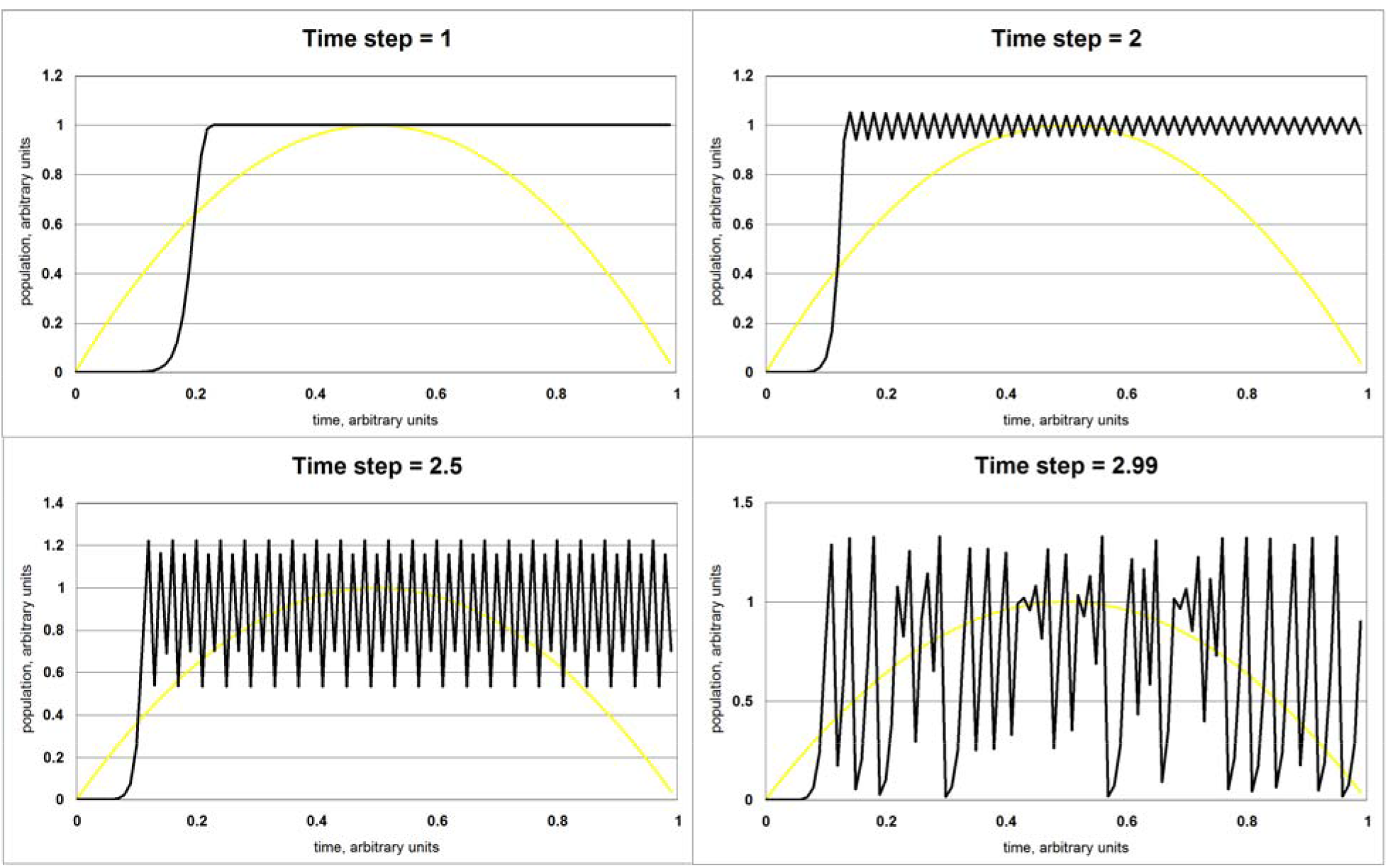
Time plot of the discrete logistic equation. As the time increment Δ t increases above ∼2.5, the behavior of the solution goes from periodic to chaotic.

## Heuristic behavior of populations with delayed logistic feedback

The value of the delay is an environmental parameter, not under genetic control. But the population growth rate depends on fertility and lifespan, which are both evolvable traits. We expect that individual selection will push fertility higher and lifespan longer, creating ever-higher growth rates at the population level. As growth rate increases, the chaos parameter approaches 3, and the local population destroys itself.

In a metapopulation, any local extinction will be temporary, as the population can be re-seeded from another population site that has not become extinct. The site that supplies founders to re-seed the extinct site is likely to have lower fertility and longevity, simply because it has not yet destroyed itself. As this process of extinction and re-seeding is iterated, the metapopulation evolves toward a growth rate near the threshold of chaos. The dynamic of extinctions and re-seeding holds the growth rate in check, so that extinctions become infrequent.

Individual selection, of course, pushes for both longer lifespans and higher fertility. A sustainable net growth rate, high enough to recover from population dips but low enough to avoid the chaos threshold, can be achieved either with low fertility or short lifespan or any combination of the two. So from the perspective of group selection, the two strategies are competitive. But reproducing quickly and dying sooner has an advantage at the level of individual selection. For the case of a stable population, it is well-known that the winning strategy is the one with higher Malthusian parameter ***r*** [36], and this corresponds to the combination of high fertility with short lifespan.

In the real world, many factors affect the optimal combination of lifespan and fertility, including incidental mortality, parental care, stability of other environmental factors, and physical constraints. We should not expect that shorter lifespans with higher fertility are always the optimal combination.

Extracted from the present computer model, Figure 2 displays two samples of population dynamics at single sites. In 2a, we see a typical progression with evolved pleiotropy. Lifespan and fertility have been initialized with low values, so population growth is slow at first compared to the delay time. As fertility and lifespan evolve higher, the dynamic becomes unstable. Starting about t=5,000, population swings are extreme and extinctions are frequent. (Populations falling to zero are immediately re-seeded, so they appear to rise again continuously in the chart.) Dramatically, around time=12,000, one of the re-seedings happens to occur by an agent evolved for stable growth rate and for awhile the population is temporarily stable, but soon individual selection pushes fertility and lifespan higher again, and the deep cycling returns. Globally, the metapopulation only survives because individual sites don’t all become extinct at once, but pockets of stability remain rare and temporary. For comparison, 2b shows a typical progression when pleiotropy is permitted to evolve. The initial march toward chaos is similar but slower. The population enters a stage of deep cycling. Then at about t=14,000, one of the re-seedings happens to occur by an agent evolved for stable growth rate. This time, the stability is not lost, because pleiotropy has evolved. The population growth rate is protected from further increase because every increase in fertility is balanced by a corresponding decrease in lifespan, and *vice versa*. Thereafter, population dynamics remains stable for many thousands of time steps (off the right edge of the graph, not pictured).

**Figure 2.**
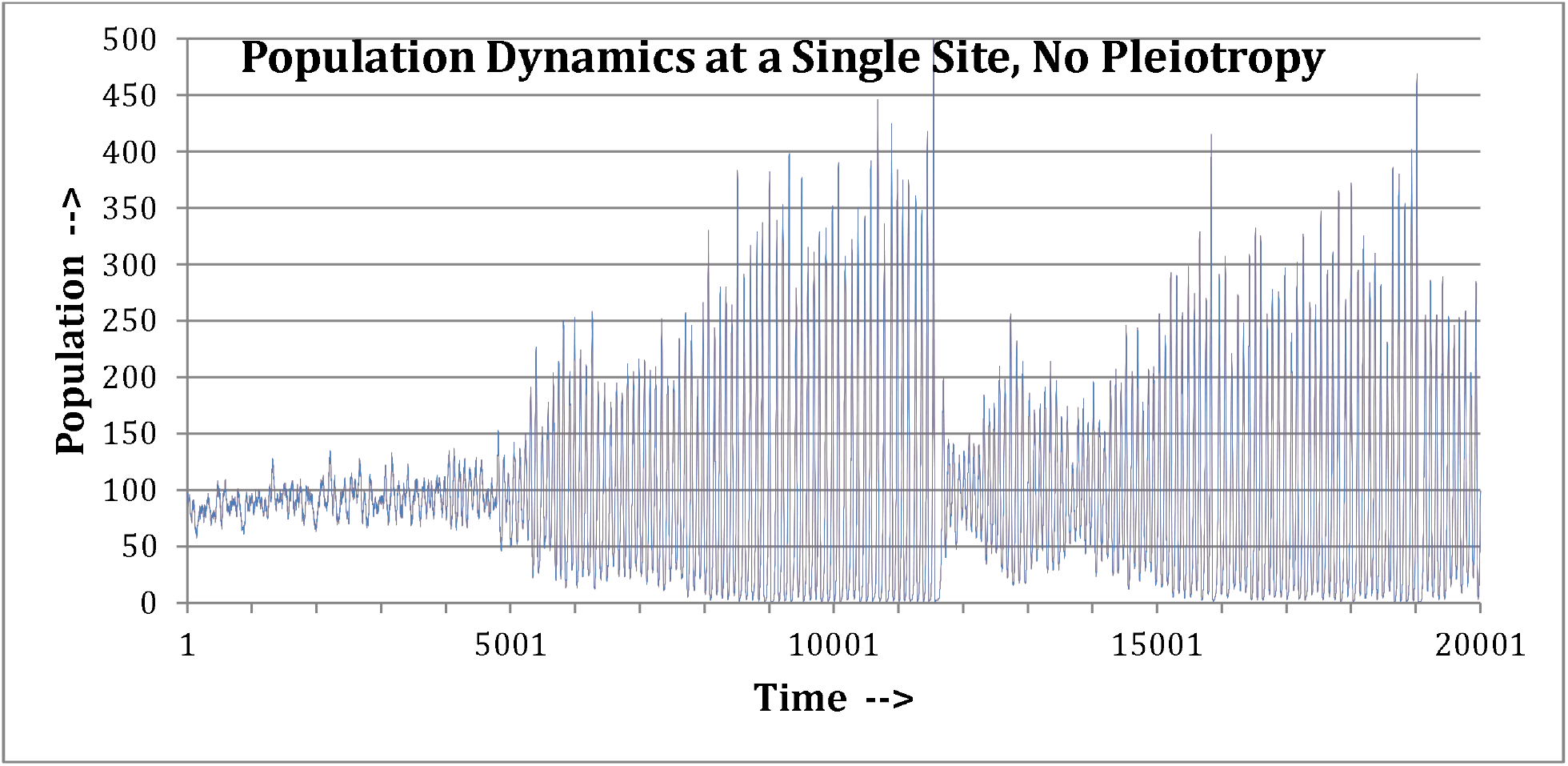
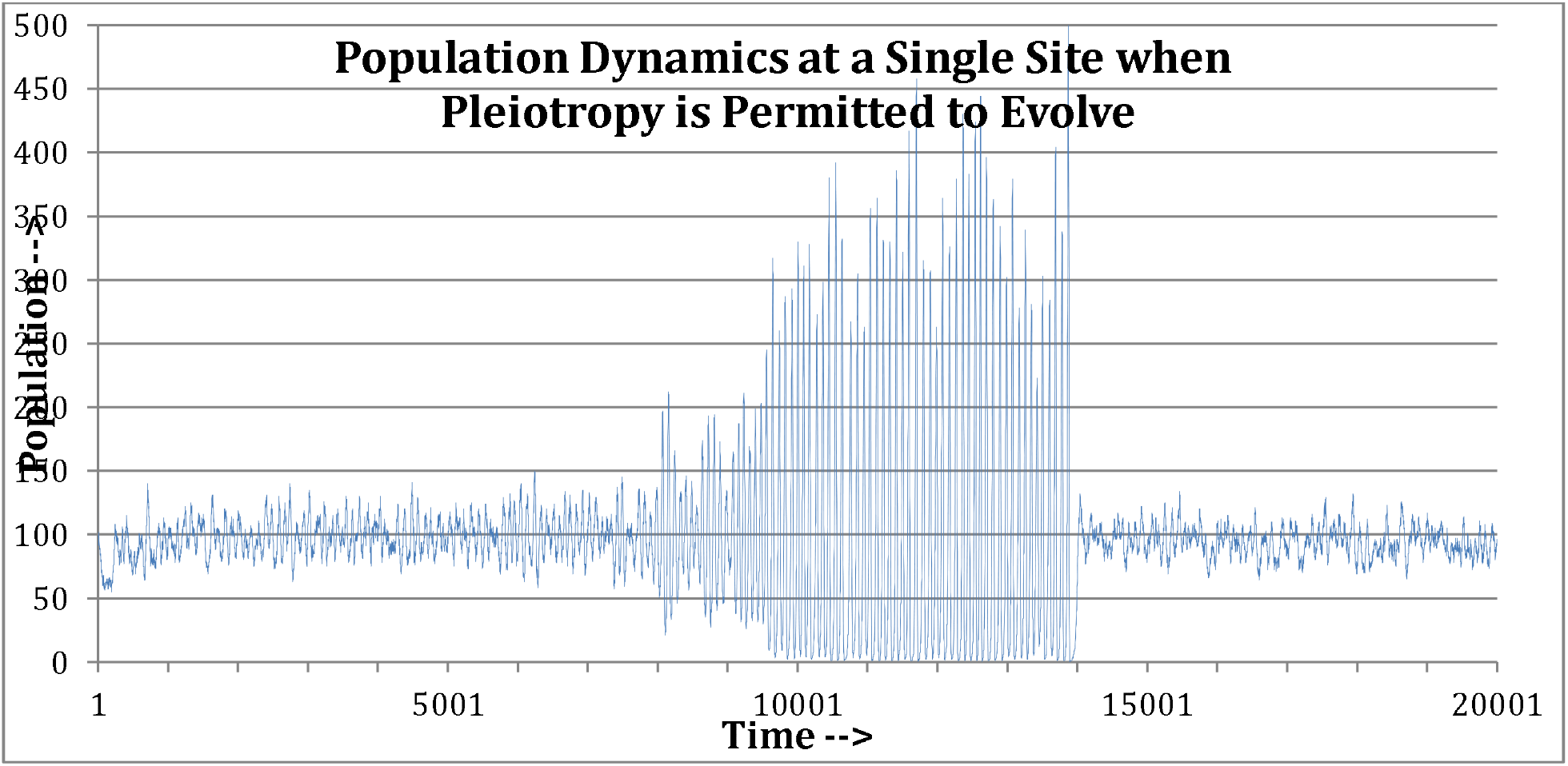
Population curves for a single subpopulation site illustrate the basic dynamics of selection within the metapopulation. (a) without pleiotropy (b) with pleiotropy.

## Metapopulation Dynamics of the Present Model

I have argued [4] and others have agreed [25] that local population dynamics can be rapid [37] and that local extinctions can be a strong and rapid selective force [38], comparable in efficiency to individual selection, but pulling in the opposite direction. This provides the basis for the most plausible scenario for the evolution of senescence. Even in the case of no pleiotropy (above, Fig. 2), growth in fertility and longevity are tempered by the discipline of continuing extinctions. Each time a population drives itself extinct, it is re-seeded from a site that is less far along on the suicidal path of accelerating growth. The re-seeding agent is likely to have lower fertility and longevity than the site’s recent (extinct) occupants. This dynamic holds lifespan and fertility permanently in check globally, albeit at levels that are just over the line of unsustainability. By itself, each site is unstable, but globally, the metapopulation can maintain itself indefinitely.

When the option of evolvable pleiotropy is added to the model, the individual sites have a resource for restraining their suicidal march toward higher growth rates. Sites that evolve a linkage between higher fertility and shorter lifespan can avoid the repeated brushes with extinction, and remain stable for a long time. If individuals in which mutations for lifespan and for fertility are inversely linked can dominate a local population, then that population can escape the hazard of continually evolving over the brink of chaos. The advantage of the stable population is realized as a greater probability that its individuals will become the founders that re-seed sites that have oscillated to extinction.

## Model Results

For a broad range of parameters, pleiotropy evolves to a value between –0.5 and –0.7. This is on a scale where 0 means that fertility and longevity mutate independently without linkage, and –1 indicates perfect inverse correlation. (The minus sign means that longer lifespan is linked to lower fertility. The model also permits positive values of pleiotropy, meaning that lifespan and fertility would rise and fall in parallel.)

Results were recorded for a range of the delay parameter, for 5 values of the migration parameter, and for two ways of selecting the founder agent that re-seeds each site after extinction, to wit: In series (A), the individual is chosen at random from the metapopulation, every individual having equal weight. In series (B), sites that are less volatile are favored, so individuals from stable sites have a better shot at re-seeding. The importance of this choice is clear, because it is only in spreading from site to site that a genotype can prevail in the metapopulation. (A) is certainly an underestimate of the force of group selection, because it gives an advantage to sites that have population blooms with many times the steady-state sustainable population. In the real world, it takes time to find a newly empty homeland ripe for colonization, and a population that blooms and dies is unlikely to export successfully. (B) is probably an overestimate of the force of group selection, because it weights too heavily in favor of the stable sites over the volatile sites which might have much larger populations to export.

For low values of migration, pleiotropy almost always evolved to a value between –0.5 and –0.7. For the highest values of migration, pleiotropy failed to evolve. This is to be expected, because sufficiently high rates of migration effectively turn a metapopulation into a unified population, and only individual selection matters.

With very high values of migration, the “profligate” strategy is optimal: the combination of long lifespan and high fertility produces high, short-lived population peaks, followed in a few cycles by local extinction. If this strategy takes over enough sites, it can sustain itself with continual re-seeding. The profligate strategy evolved with positive values of the pleiotropy parameter, the opposite of antagonistic pleiotropy. In option (A) the profligate strategy did tolerably well for high values of the delay parameter, but appeared only rarely for low values. In option (B), the profligate parameter almost always fared poorly. Negative pleiotropy (AP) evolved in over 95% of the runs, but occasionally there was an outlier run in which the profligate strategy took over.

Figure 3 displays the low-migration results. 3(A) is re-seeding with a random agent (option (A)). 3(B) is reseeding with an algorithm that gives more weight to low-volatility sites. Notice that both results stay almost always in the domain of evolved negative pleiotropy; but for option (A), “low migration” means 0 or 1×10^−4^. In option (A) the runs with migration=3×10^−4^ and 6×10^−4^ both evolve pleiotropy at low values of delay, then cross over to no pleiotropy or positive pleiotropy for higher values of delay, whereas for option (B) “low migration” behavior continues through 6×10^−4^, with a few outliers only at the 6×10^−4^ level.

**Figure 3.**
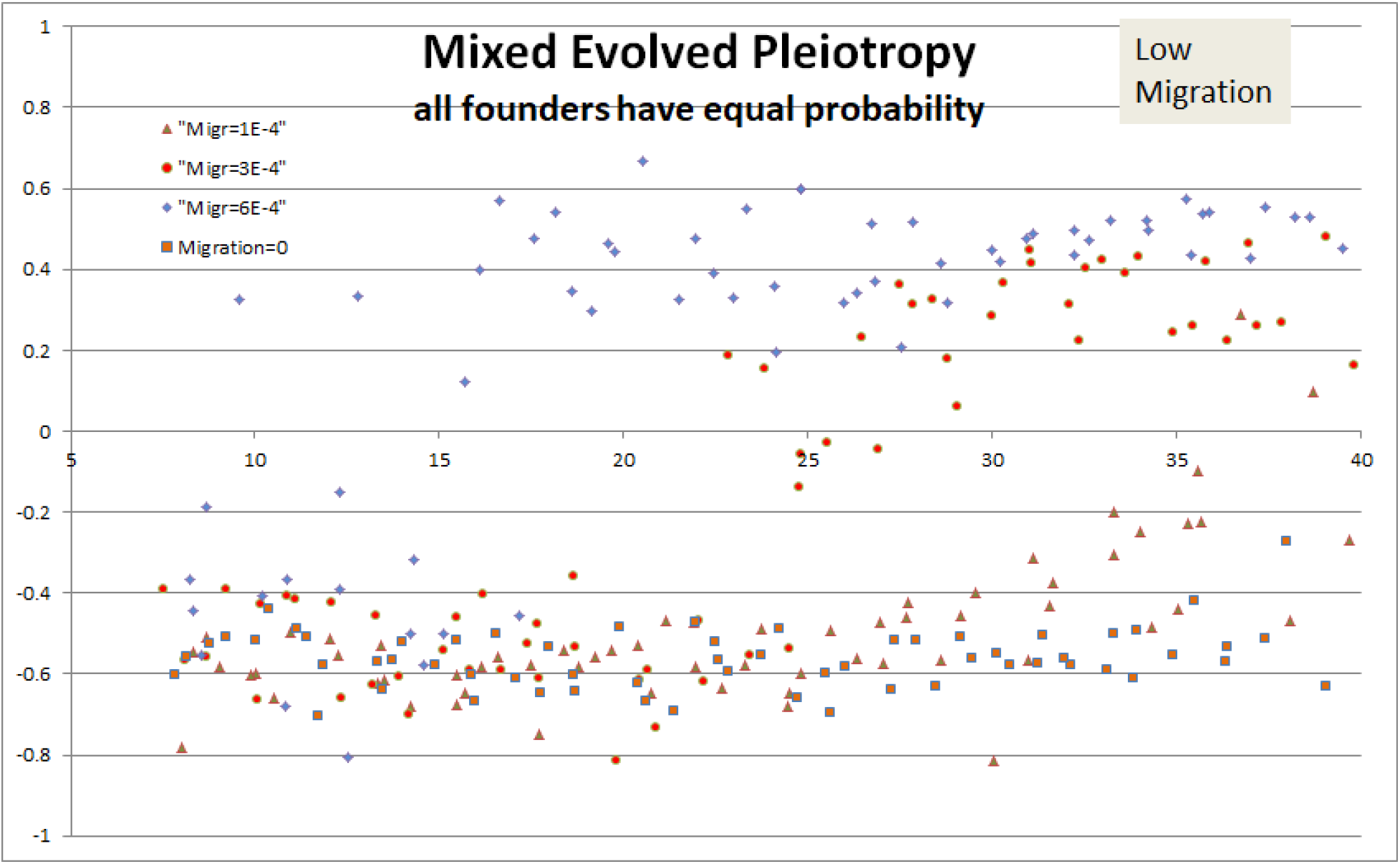
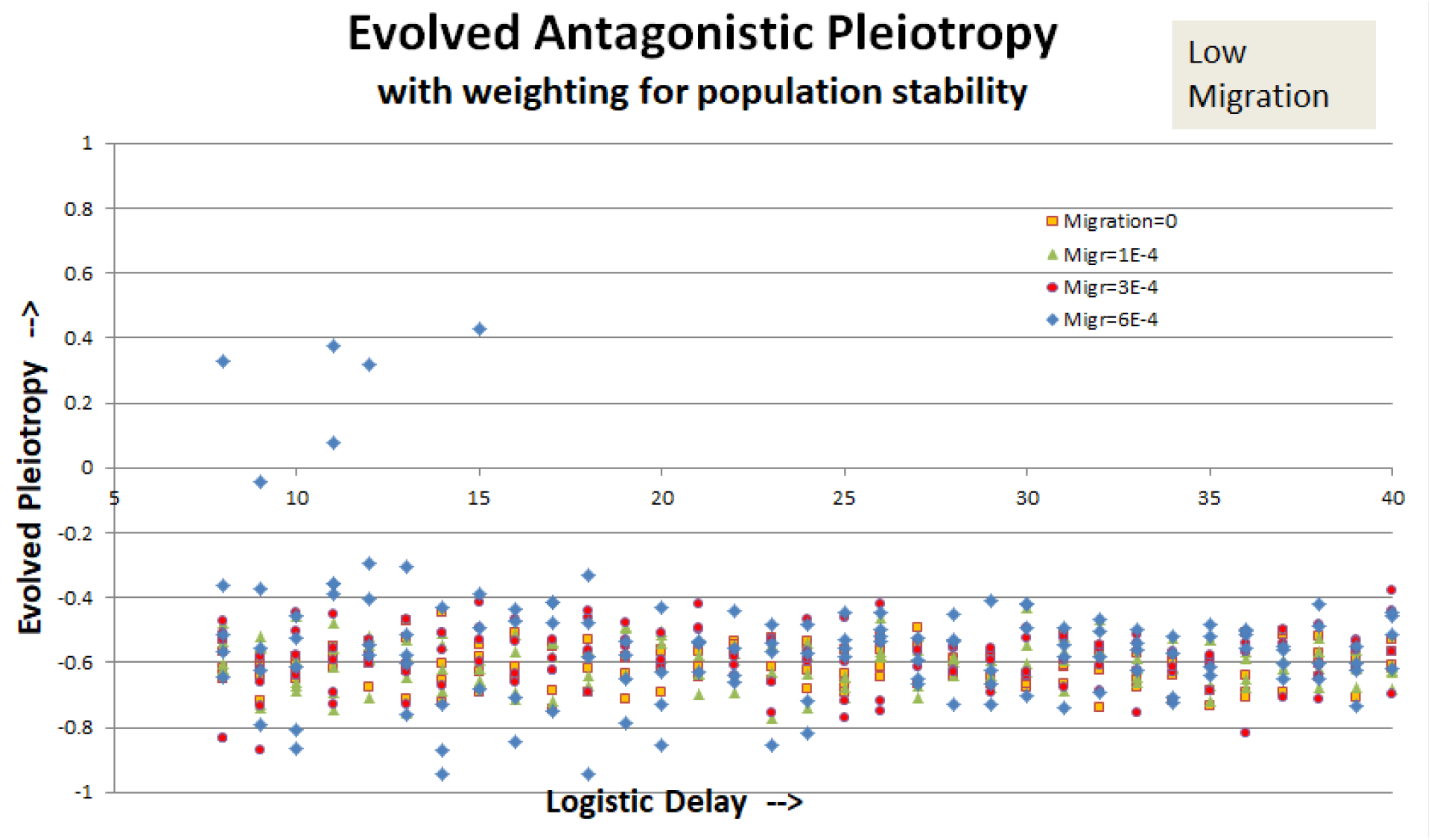
Results for evolved pleiotropy with low migration rates. (A) Every agent has equal opportunity to emigrate. (B) Preference is given to more stable sites in emigration and re-seeding.

Figure 4 displays the high-migration results. Option (A) does not evolve antagonistic pleiotropy. Option (B) continues to evolve antagonistic pleiotropy most of the time at migration=9×10^−4^ and about half the time even at 12×10^−4^.

**Figure 4.**
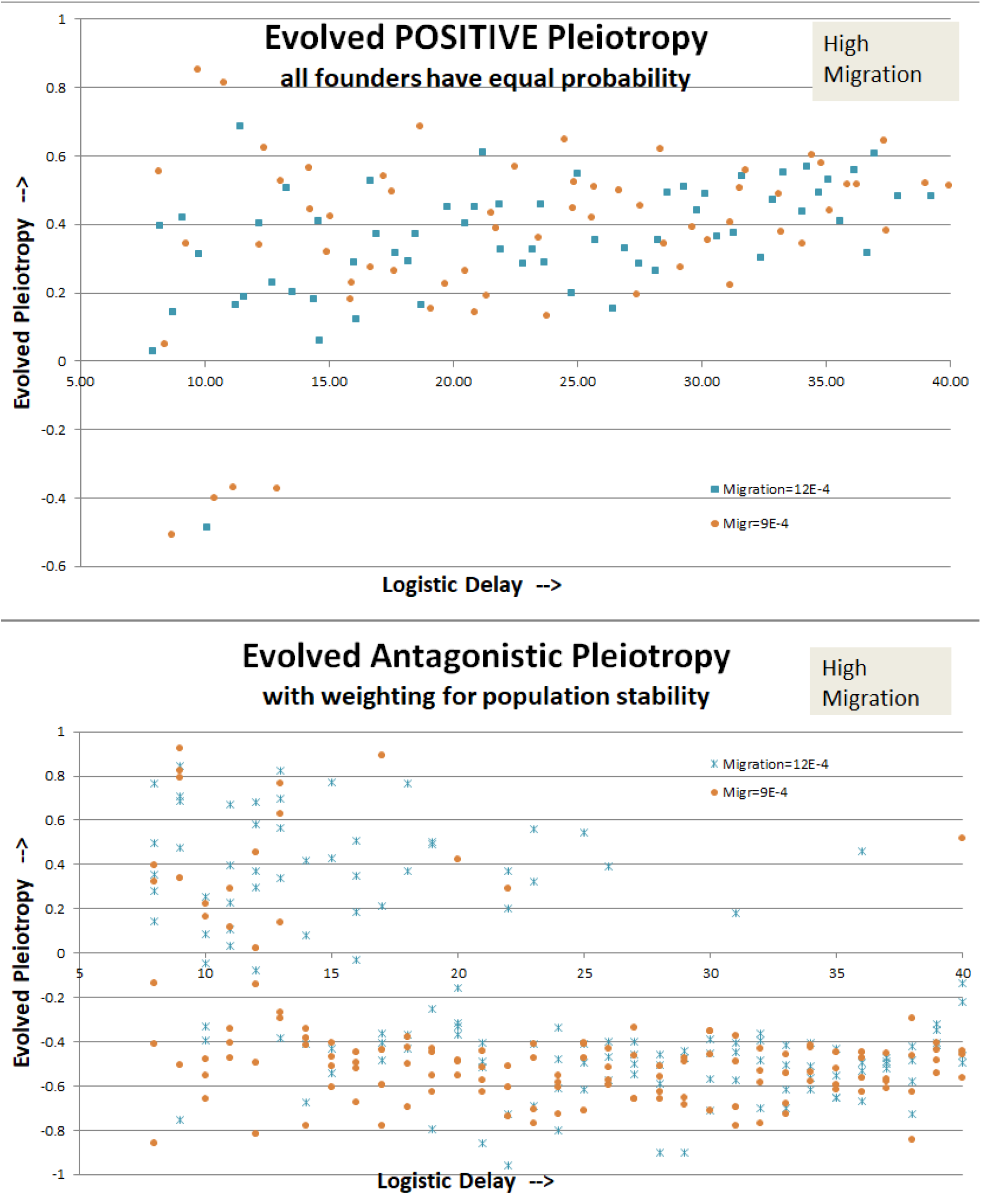
Results for evolved pleiotropy with high migration rates. (A) Every agent has equal opportunity to emigrate. (B) Preference is given to more stable sites in emigration and re-seeding.

## Detailed Description of the Model

The individual-based model consists of 100 sites, each governed by delayed logistic population dynamics with a steady-state population of 100 agents. The logistic delay (measured in time steps) is a parameter of the model. Agents have 3 evolving genes:

- Gene for lifespan. Senescence is a step function, and the agent dies abruptly when its age reaches the genetically-imposed limit.
- Gene for fertility, the probability (0<p<1) of creating a (mutated) clonal copy in each time step.
- Gene for pleiotropy. This is a number between –1 and 1 which affects the mutations of fertility and lifespan genes. For pleiotropy=0, fertility and lifespan mutations are independent. For pleiotropy=±1, fertility and lifespan mutate in lockstep, with higher fertility always leading to proportionally longer (shorter) lifespan. For intermediate values of pleiotropy, there is a proportional mix of linked mutations and independent mutations.

In each time step,

- Each agent may reproduce clonally with a probability determined by its fertility gene. Offspring genes undergo small, random and linked mutations.
- Each agent must die of old age if its age exceeds its lifespan gene.
- Each agent may die of crowding, with a probability that is proportional to the site population as it was D time steps in the past, where D=delay is a parameter of the model. (Specifically, the probability is [delayed pop]/100 * [avg fertility]. 100 is the steady-state population, always the same for all sites and times. Avg fertility is the site average of all the agents at that site. This formula assures that for each site, time-average birth rate = death rate when population = 100.

When a site oscillates to extinction, it is reseeded by a single random agent from one of the remaining sites (A). In a variation, the founding agent is chosen with a weighting factor for site life; sites that have lasted longer without oscillating to extinction are favored (B). The justification for this feature is that for the sake of computational efficiency we are re-seeding the vacant site immediately, but in reality the process is stochastic and requires time, favoring those sites that last for a time.

Optionally, there is a probability (for each site in each time step) for random emigration from another site. The emigrant agent may be chosen at random or, as above, it may be chosen with a weighting factor that prefers sites that have lasted longer. The migration rate is one parameter affecting the relative strengths of group- and individual-selection.

Each run was initialized for 25,000 time steps to allow fertility, lifespan, and pleiotropy to evolve past the point where initial conditions have been lost from the system’s memory. (This is a complex system with unpredictable behaviors, so it does not reliably reach a steady state in this time frame, or ever.) After 25,000 time steps, the system begins averaging on three different time scales. After the three averages converge to agreement, generally 40,000–50,000 time steps, the run is ended and results recorded.

## Discussion

When Williams [1] proposed the AP theory of aging, it was 1957, and his concept of genetics was Mendel’s. It was natural to assume, by analogy with human machines, that genes generally corresponded to traits, and that multiple functions for one gene was an exception. We now know that biological organisms are not designed the way humans would design them, and that multiple overlapping functions for a gene are the rule, not the exception. We should not be surprised if genes that regulate lifespan have other functions because most genes have more than one function. What is surprising is that despite the high cost of senescence in the wild, evolution has not been able to bypass pleiotropy by separating the benefit from the cost. In every other component of fitness, a strength in one area does not entail a necessary cost in another. (We don’t expect that animals that can fly must have compromised eyesight, or that animals with a keen sense of smell must pay a price with diminished speed or strength.) I suggest that there may be an adaptive value to maintaining an antagonistic link between higher fertility and shorter lifespan.

Traditionally, every time pleiotropy is observed in connection with a gene that affects lifespan, it has been interpreted as vindication of Williams and corroboration of the pleiotropic theory. However, we have seen that pleiotropy can be imposed as a precondition or it can be evolved after the fact. It can be mandatory or it can be optional. Williams’s theory applies to imposed, mandatory pleiotropy. If pleiotropy evolves—if pleiotropy derives from natural selection for pleiotropy—then it has a very different meaning. Observations of evolved pleiotropy should not be interpreted as support for the AP theories of aging, but rather as support for programmed theories, in which aging has adaptive value at the population level.

In the real world, there is sometimes pleiotropy, sometimes not, agreeing with our results. In some of our runs, the profligate strategy wins. In the real world, profligacy is limited to microbes, with rare but spectacular exceptions, like locust plagues [39], rabbits in Australia [40], and other invasive species [41].

When we observe pleiotropy, how can we tell whether it is imposed or evolved? I have argued that imposed pleiotropy is inescapable. It is a beneficial trait that logically implies long-term degradation. Some frequently-cited examples of pleiotropy in humans include the following:

- Some genes are important during development, but they fail to be turned off or turned down late in life, with consequences that accelerate aging. These include MTOR [42] and several steroid hormones. Most infamous is FSH, which rises after menopause, with no apparent purpose except to increase risk of heart attack in human females [43]. All these factors, being under epigenetic control, are examples of avoidable pleiotropy.
- Insulin/IGF signaling is necessary for growth and development, but it also increases cancer risk and shortens lifespan [44]. Like TOR, this seems to be better explained by epigenetic programming than by as an unavoidable tradeoff.
- Inflammation is a first line of defense against microbes and a core means of degrading damaged tissue. Nevertheless, inflammaging is the best-studied mechanism that accelerates aging and increases risk of all the diseases of old. Inflammaging may be interpreted as a pleiotropic consequence of inflammation [45], but inflammation contributes in no detectable way to aging during the first decades of life. Thus it might also be seen as the body recruiting a mechanism of self-defense for an application of self-destruction late in life.
- Recently, methylation patterns associated with accelerated aging have been associated with the expression of telomerase, which helps maintain cells in a youthful state [46]. There seems to be no logical reason why these two functions should be connected, so it is a candidate to be understood as evolved pleiotropy.

These examples and others are reviewed by Leroi et al [5], who ask why so few examples of AP have been identified in animal biology but do not question the fundamental validity of the theory that aging is caused by unavoidable tradeoffs.

Imposed antagonistic pleiotropy is an explanation for why aging evolved despite its individual cost. Evolved antagonistic pleiotropy is a protective trait, a group-selected adaptation built atop the earlier group-selected adaptation of senescence itself.

